# MSnbase, efficient and elegant R-based processing and visualisation of raw mass spectrometry data

**DOI:** 10.1101/2020.04.29.067868

**Authors:** Laurent Gatto, Sebastian Gibb, Johannes Rainer

## Abstract

We present version 2 of the `MSnbase` R/Bioconductor package. `MSnbase` provides infrastructure for the manipulation, processing and visualisation of mass spectrometry data. We focus on the new *on-disk* infrastructure, that allows the handling of large raw mass spectrometry experiments on commodity hardware and illustrate how the package is used for elegant data processing, method development, and visualisation.

## Introduction

Mass spectrometry is a powerful technology to assay chemical and biological samples. It is used in routine applications with well characterised protocols such as in clinical settings, as well as a development platform, with the aim to improve on existing protocols and devise new ones. The complexity and diversity of mass spectrometry yield complex data of considerable size, that require non trivial processing before producing interpretable results. The complexity and size of these data constitute a significant challenge for protocol development: in addition to the development of sample processing and mass spectrometry methods that yield the raw data, it is essential to process, analyse, interpret and assess these new data to demonstrate the improvement in the technical, analytical and computational workflows.

Practitioners have a diverse catalogue of software tools at their disposal. These range from low level software libraries that are aimed at programmers to enable the development of new applications, to more user-oriented applications with graphical user interfaces which provide a more limited set of functionalities to address a defined scope. Examples of software libraries include Java-based jmzML^1^ or C/C++-based ProteoWizard.^2^ Thermo Scientific Proteome Discoverer (Thermo Fisher Scientific), MaxQuant^3^ and PeptideShaker^4^ are among the most widely used user-centric applications.

In this software note, we present version 2 of the `MSnbase`^5^ software, available from the Bioconductor^6^ project. The package, like other software such as Python-based pyOpenMS,^7^ spectrum_utils or Pyteomics, offers a platform that lies between low level libraries and end-user software. `MSnbase` provides a flexible R^l0^ command-line environment for metabolomics and proteomics mass spectrometry-based applications. It lays out a sound infrastructure to work with raw mass spectrometry data from MS files in mzML, mzXML, mzData or ANDI-MS/netCDF format as well as quantitative and proteomics identification data. The package enables manipulation (for example subsetting, filtering, or accessing specific parts thereof), detailed step-by-step processing (for example smoothing and centroiding of profile-mode MS data, or normalisation and imputation of quantitative data), analysis and visualisation of these data and the development of novel computational mass spectrometry methods.^11^ Extensive documentation and use cases are provided in *package vignettes*^l2^ and workflows.^l3^ Here, we focus on the new developments pertaining to raw mass spectrometry data handling and processing.

### Infrastructure for raw data

In `MSnbase`, mass spectrometry experiments are handled as `MSnExp` objects. While the implementation is more complex, it is useful to schematise a raw data experiment as being composed of raw data, i.e. a collection of individual spectra, as well as spectra-level metadata (Figure 1). Each spectrum is composed of m/z values and associated intensities. The metadata are represented by a single table with variables along the columns and each row associated to a spectrum. Among the metadata available for each spectrum, there are MS level, acquisition number, retention time, precursor m/z and intensity (for MS level 2 and above), and many more. `MSnbase` relies on the `mzR` package^2^ to import raw mass spectrometry data from one of the many community-maintained open standards formats (mzML, mzXML, mzData or ANDI-MS/netCDF) and provides a rich and principled interface to manipulate such objects. The code chunk below illustrates such an object as displayed in the R console and an enumeration of the metadata fields.

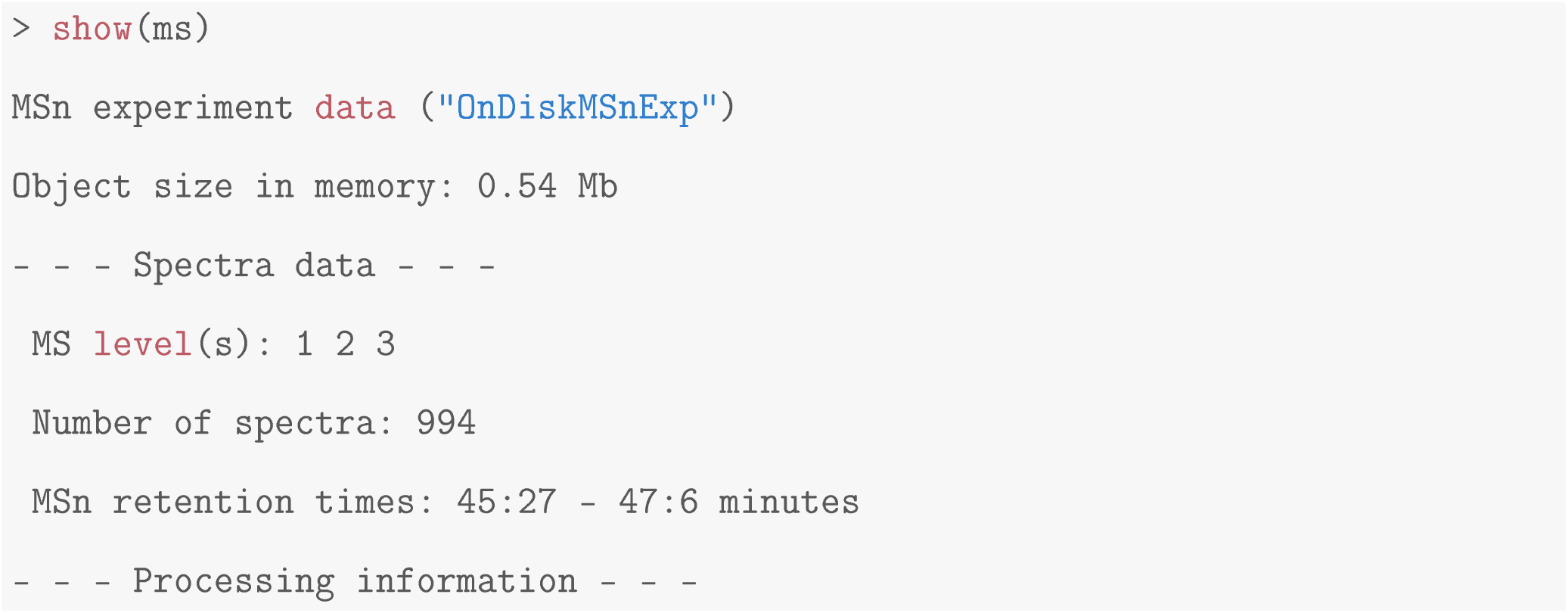

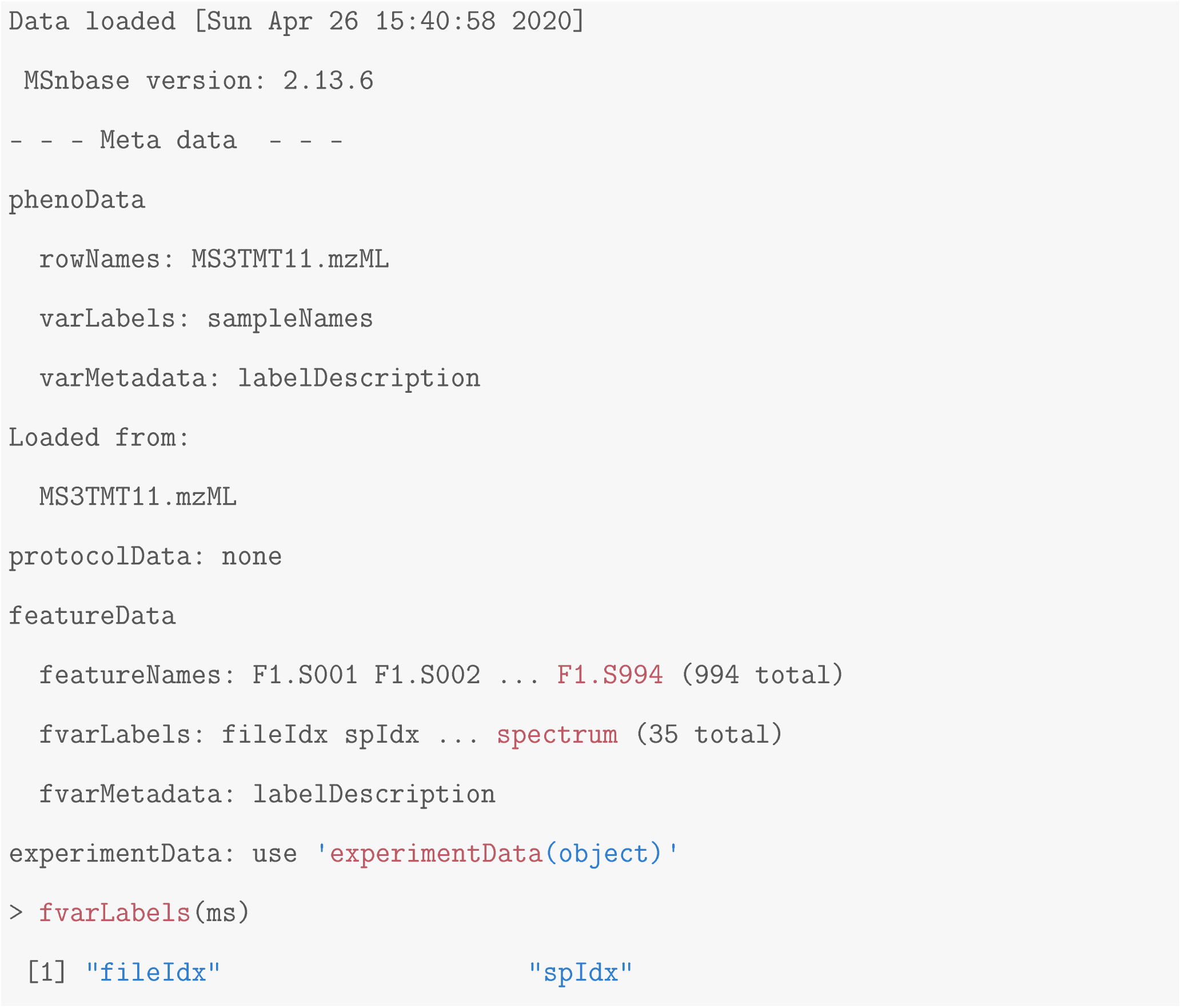

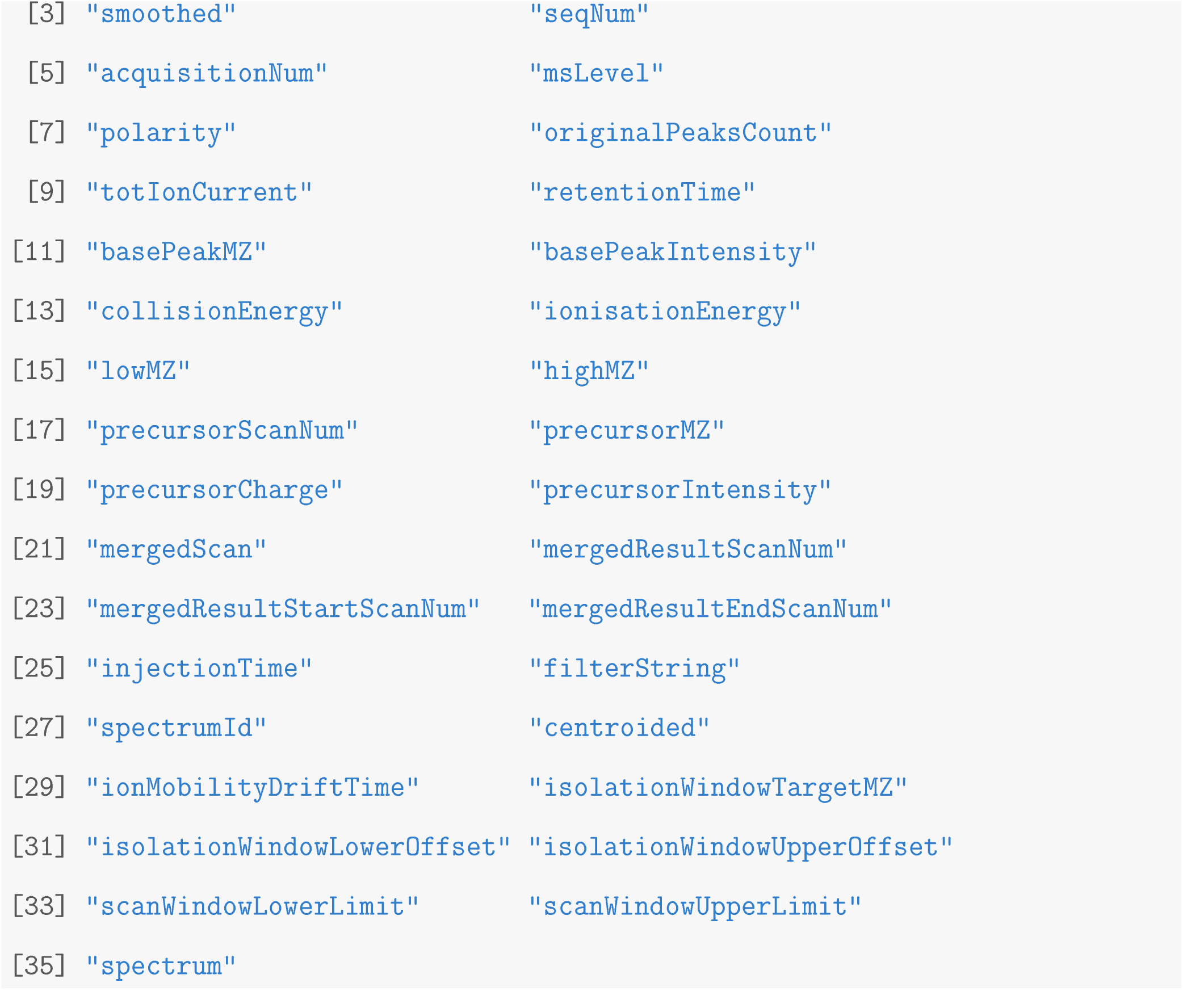

**Figure 1:**
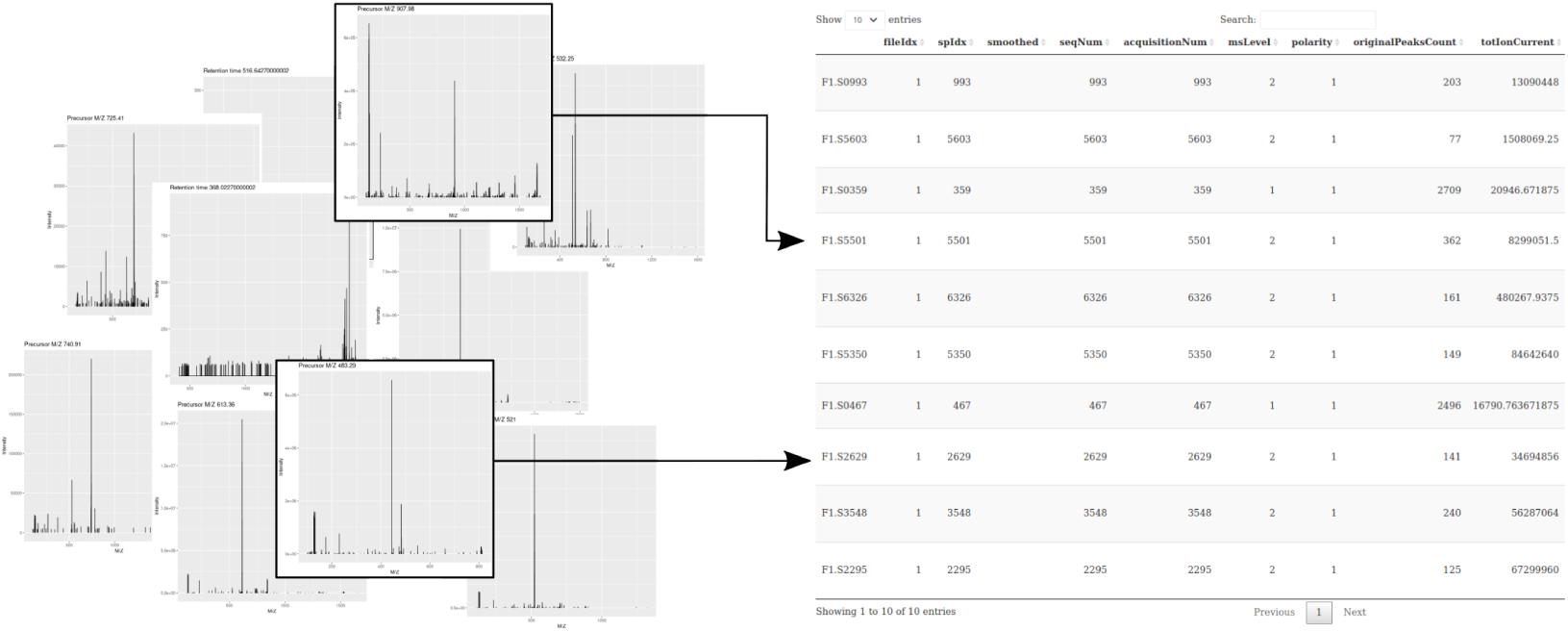
Schematic representation of what is referred to by *raw data*: a collection of mass spectra and a table containing spectrum-level annotations along the lines. Raw data are imported from one of the many community-maintained open standards formats mzML, mzXML, mzData or ANDI-MS/netCDF).

In the following sections, we describe how `MSnbase` can be used for data processing and visualisation. An example of its ability to also efficiently handle very large mass spectrometry experiments (in this case with 3,773,464 spectra in 1,182 mzXML files) is provided as supplementary information. We will also illustrate how it makes use of the forward-pipe operator (`%>%`) defined in the `magrittr` package. This operator has proved useful to develop non-trivial analyses by combining individual functions into easily readable and elegant pipelines.

### On-disk backend

The main feature in version 2 of the `MSnbase` package was the addition of different backends for raw data storage, namely *in*-*memory* and *on-disk*. The following code chunk demonstrates how to import data from an mzML file to create two `MSnExp` objects that store the data either in memory or on disk.

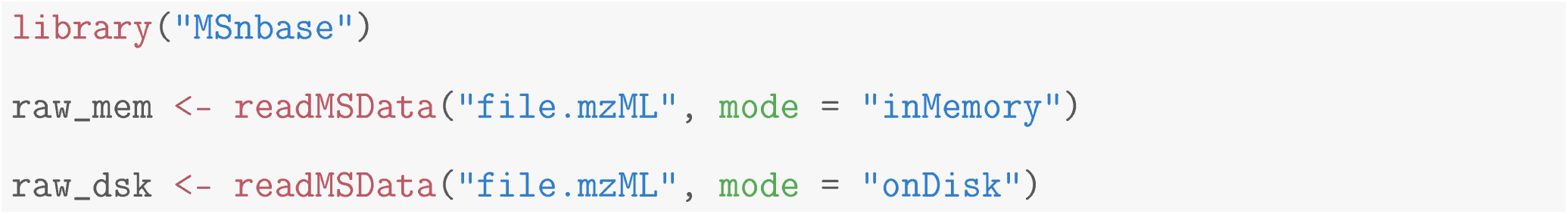

Both modes rely on the `mzR` ^2^ package to access the spectra (using the `mzR ::peaks()` function) and the metadata (using the `mzR ::header()` function) in the data files. The former is the legacy storage mode, implemented in the first version of the package, that loads all the raw data and the metadata into memory upon creation of the in-memory `MSnExp` object. This solution doesn’t scale for modern large dataset, and was complemented by the on-disk backend. The on-disk backend only loads the metadata into memory when the on-disk `MSnExp` is created and accesses the spectra data (i.e. m/z and intensity values) in the original files on disk only when needed (see below and Figure 2 (d)), such as for example for plotting. There are two direct benefits using the on-disk backend, namely faster reading and reduced memory footprint. Figure 2 shows 3-fold faster reading times (a) and over a 10-fold reduction in memory usage (b).

**Figure 2:**
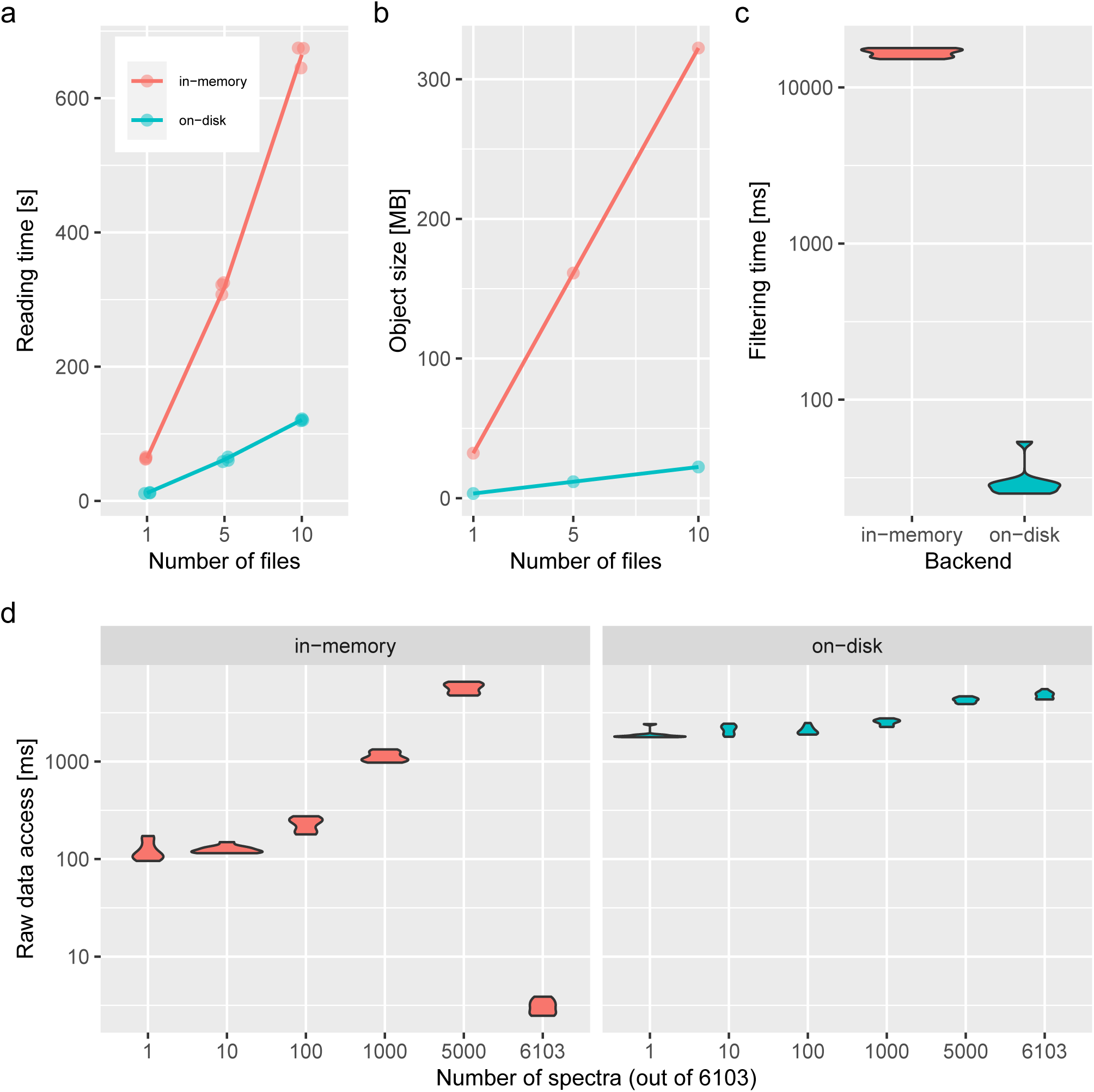
(a) Reading time (triplicates, in seconds) and (b) data size in memory (in MB) to read/store 1, 3 and 10 files containing 1431 MS1 (on-disk only) and 6103 MS2 (on-disk and in-memory) spectra. (c) Filtering benchmark assessed over 10 interactions on in-memory and on-disk data containing 6103 MS2 spectra. (d) Access time to spectra for the in-memory (left) and on-disk (right) backends for 1, 10, 100 1000, 5000 and all 6103 spectra. Benchmarks were performed on a Dell XPS laptop with an Intel i5-8250U processor 1.60 GHz 4 cores, s threads), 7.3 GB RAM running Ubuntu 18.04.4 LTS 64-bit and an SSD drive. The data used for the benchmarking are a TMT 4-plex experiment acquired on a LTQ Orbitrap Velos (Thermo Fisher Scientific) available in the `msdata` package and described in^l4^.

Because the on-disk backend does not hold all the spectra data in memory, direct manipulations of these data are not possible. We thus implemented a t mechanism for this backend that caches any data manipulation operations in a processing queue in the object itself. These operations are then applied only when the user accesses m/z or intensity values. As an additional advantage, operations on subsets of the data become much faster since data manipulations are applied only to data subsets instead of the full data set at once. Also, on-disk data access is parallelized by data file ensuring a higher performance of this backend over conventional in-memory data representations. As an example, the following short analysis pipeline, that can equally be applied to in-memory or on-disk data, retains MS2 spectra acquired between 1000 and 3000 seconds, extracts the m/z range corresponding to the TMT 6-plex range and focuses on the MS2 spectra with a precursor intensity greater than 11 *×* 10^6^ (the median precursor intensity).

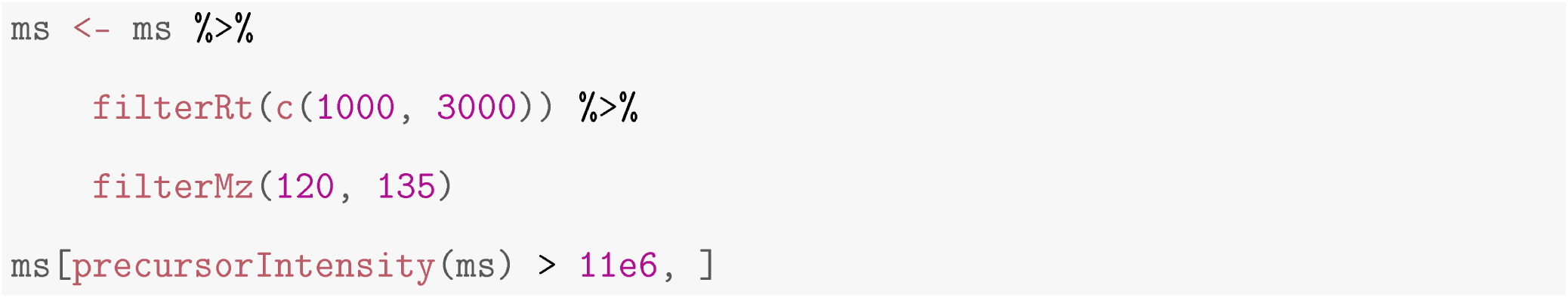

As shown on Figure 2 (c), this lazy mechanism is significantly faster than its application on in-memory data. The advantageous reading and execution times and memory footprint of the on-disk backend are possible by retrieving only spectra data from the selected subset hence avoiding access to the full raw data. Once access to the spectra m/z and intensity values become mandatory (for example for plotting), then the in-memory backend becomes more efficient, as illustrated on Figure 2 (d). The benefit of accessing data in memory is however reduced by underlying copies that are performed during the subsetting operation. When subsetting an in-memory `MSnExp` into a new, smaller in-memory `MSnExp` instance, the matrices that contain the spectra for the new object are copied, thus leading to increased execution time and (transient, if the original data are replaced) memory usage. Figure 2 (d) shows that the larger the subset, the smaller the benefits of an in-memory backend become. The example with the 6103 spectra, corresponding to the full data (i.e. all spectra are already in memory and there is no memory management overhead) is representative of memory access only and constitutes the best case scenario.

The on-disk backend has become the preferred backend for large data, and the only viable alternative when the size of the data exceeds the available RAM and/or when several MS levels are to be loaded and handled simultaneously. The in-memory backend can still prove useful in cases when small MS2-only data are to be analysed, and will remain available in future versions of `MSnbase`.

### Prototyping

The `MSnExp` data structure and its interface constitute an efficient prototyping environment for computational method development. We illustrate this by demonstrating how to implement the BoxCar^l5^ acquisition method. In a nutshell, BoxCar acquisition aims at improving the detection of intact precursor ions by distributing the charge capacity over multiple narrow m/z segments and thus limiting the proportion of highly abundant precursors in each segment. A full scan is reconstructed by combining the respective adjacent segments of the BoxCar acquisitions. The `MSnbaseBoxCar` package^l6^ is a small package that demonstrates this. The simple pipeline is composed of three steps, described below, and illustrated with code from `MSnbaseBoxCar` in the following code chunk.

1. Identify and filter the groups of spectra that represent adjacent BoxCar acquisitions (Figure 3 (b)). This can be done using the *filterString* metadata variable that identifies BoxCar spectra by their adjacent m/z segments with the `bc_groups()`function and filtering relevant spectra with the `filterBoxCar()`.
2. Remove any signal outside the BoxCar segments using the `bc_zero_out_box()` function from `MSnbaseBoxCar` (Figures 3 (c) and (d)).
3. Using the `combineSpectra` function from the `MSnbase`, combine the cleaned BoxCar spectra into a new, full spectrum (Figure 3 (e)).

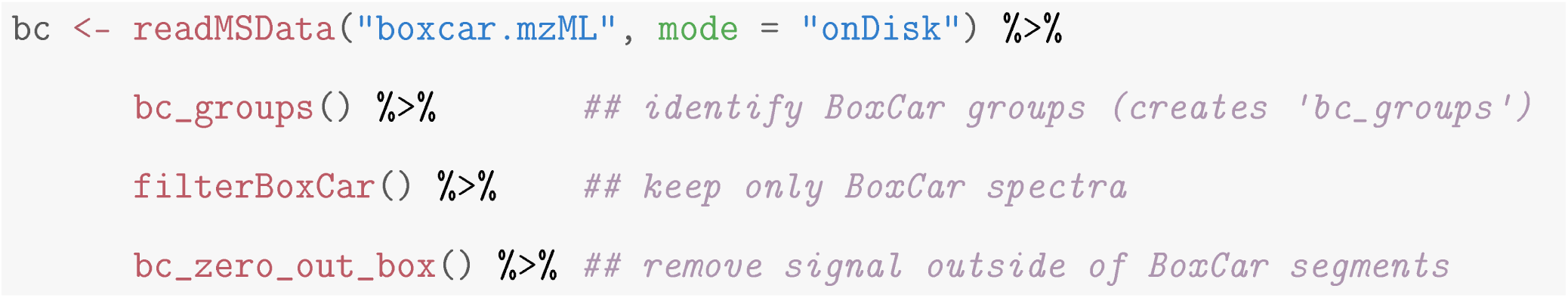

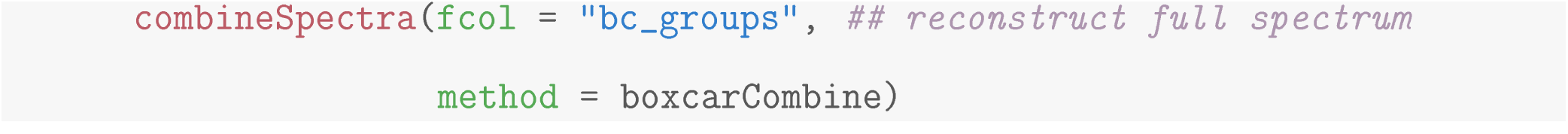

**Figure 3:**
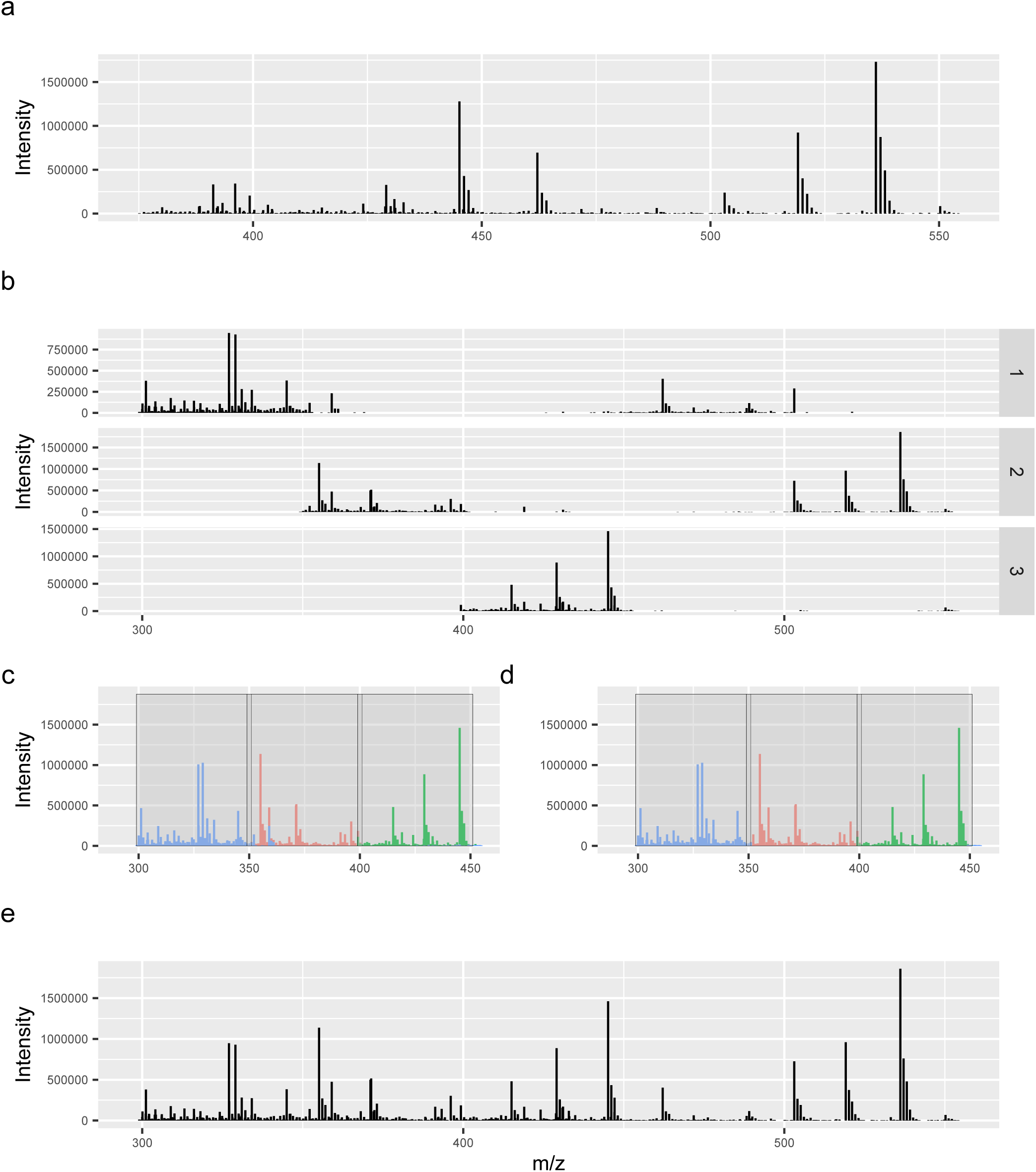
BoxCar processing with `MSnbase`. (a) Standard full scan with (b) three corresponding BoxCar scans showing the adjacent segments. Figure (c) shows the overlapping intact BoxCar segments and (d) the same segments after cleaning, i.e. where peaks outside of the segments were removed. The reconstructed full scan is shown on panel (e). Spectra visualisation, as shown here, rely on the ggplot2^l7^ package.

After processing of the BoxCar data, the final object can either be further analysed using `MSnbase` or written back to disk as an `mzML` file using `writeMSData()` for processing with other tools.

All the functions for the processing of BoxCar spectra and segments in `MSnbaseBoxCar` were developed using existing functionality implemented in `MSnbase`, illustrating the flexibility and adaptability of the `MSnbase` package for computational mass spectrometry method development.

### Visualisation

The R environment is well known for the quality of its visualisation capacity. This also holds true for mass spectrometry.^l8 -2l^ Here, we conclude the overview of version 2 of the `MSnbase` package by highlighting the flexibility of the software to visualise and assess the efficiency of raw data processing. Figure 4 compares the raw MS profile data imported from an mzML file for serine and the same data after smoothing, centroiding and m/z refinement, as illustrated in the code chunk below. Detailed execution and description of these operations can be found in the *Msnbase: centroiding of profile-mode MS data* `MSnbase` vignette.

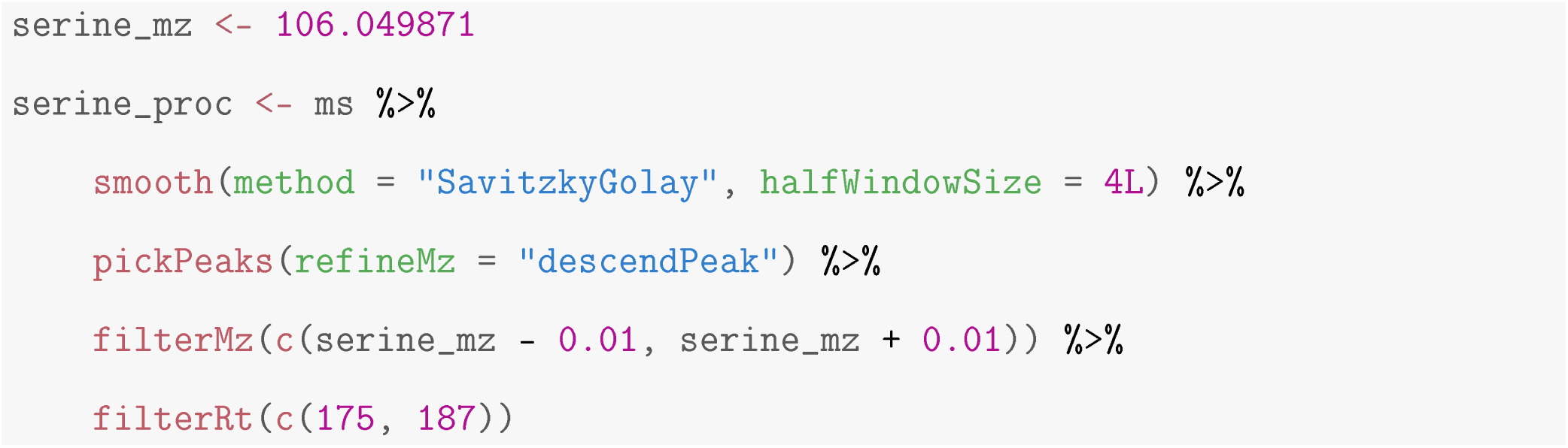

**Figure 4:**
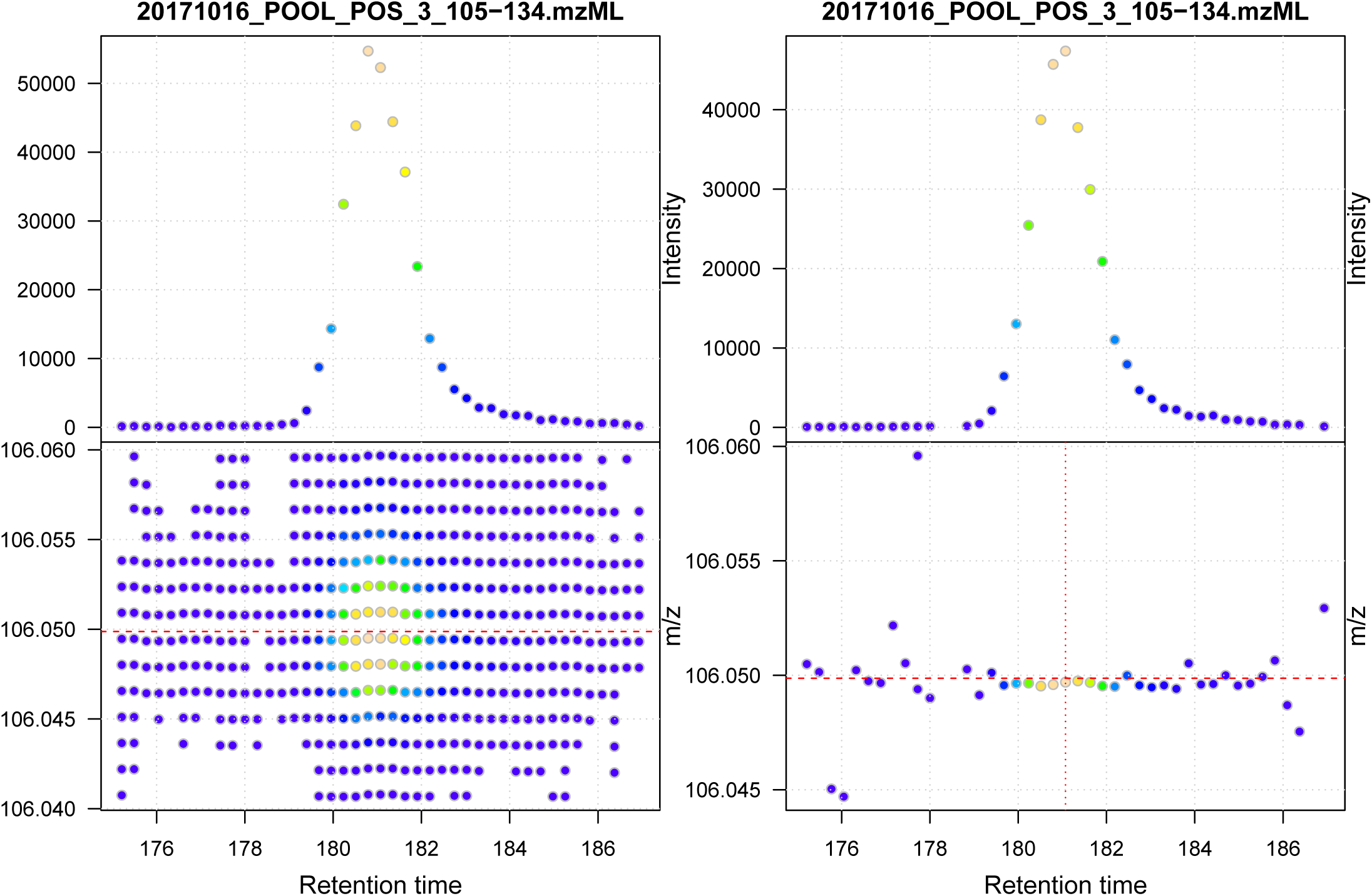
Visualisation of data smoothing and m/z refinement using `MSnbase`. (a) Raw MS profile data for serine. Upper panel shows the base peak chromatogram (BPC), lower panel the individual signals in the retention time - m/z space. The horizontal dashed red line indicates the theoretical m/z of the [M+H]+ adduct of serine. (b) Smoothed and centroided data for serine with m/z refinement. The horizontal red dashed line indicates the theoretical m/z for the [M+H]+ ion and the vertical red dotted line the position of the maximum signal.

### Package maintenance and governance

The first public commit to the `MSnbase` GitHub repository was in October 2010. Since then, the package benefited from 12 contributors^22^ that added various features, some particularly significant ones such as the on-disk backend described herein. Contributions to the package are explicitly encouraged, rewarded by an official contributor status and governed by a code of conduct.

According to `Msnbase’` s Bioconductor page, there are 36 Bioconductor packages that depend, import or suggest it. Among these are pRoloc^23^ to analyse mass spectrometry-based spatial proteomics data, msmsTests,^24^ DEP,^25^ DAPAR and ProStaR ^26^ for the statistical analysis of quantitative proteomics data, RMassBank^27^ to process metabolomics tandem MS files and build MassBank records, MSstatsQC^2^ for longitudinal system suitability monitoring and quality control of targeted proteomic experiments and the widely used xcms^2^ package for the processing and analysis of metabolomics data. `MSnbase` is also used in non-R/Bioconductor software, such as for example IsoProt,^30^ that provides a reproducible workflow for iTRAQ/TMT experiments. The BioContainers^3l^ project overs a dedicated container for the `MSnbase` package, this facilitating the reuse of the package in third-party pipelines. `MSnbase` currently ranks 101 out of 1823 packages based on the monthly downloads from unique IP addresses, tallying over 1000 downloads from unique IP addresses each months.

As is custom with Bioconductor packages, `MSnbase` comes with ample documentation. Every user-accessible function is documented in a dedicated manual page. In addition, the package offers 3 vignettes, including one aimed at developers. The package is checked nightly on the Bioconductor servers: it implements unit tests covering 72% of the code base and, through its vignettes, also provides integration testing. Questions from users and developers are answered on the Bioconductor support forum as well as on the package GitHub page. The package provides several sample and benchmarking datasets, and relies on other dedicated *experiment packages* such as msdata^32^ for raw data or pRolocdata^23^ for quantitative data. `MSnbase` is available on Windows, Mac OS and Linux under the open source Artistic 2.0 license and easily installable using standard installation procedures.

The growth of `MSnbase` and the user support provided over the years attest to the core maintainers commitment to long-term development, and the quality and maintainability of the code base.

## Discussion

We have presented here some important functionality of `MSnbase` version 2. The new on-disk infrastructure enables large scale data analyses,^33^ either using `MSnbase` directly or through packages that rely on it, such as `xcms.` We have also illustrated how `MSnbase` can be used for standard data analysis and visualisation, and how it can be used for method development and prototyping.

The version of `MSnbase` used in this manuscript is 2.14.2. The main features presented here were available since version 2.0. The code to reproduce the analyses and figures in this article is available at https://github.com/lgatto/2020-msnbase-v2/.

## Associated Content

Supplementary file 1: script documenting the processing of 1182 mzXML files (5773464 spectra) using MSnbase.

## Acknowledgenent

The authors thank the various contributors and users who have provided constructive input and feedback that have helped, over the years, the improvement of the package. The authors declare no conflict of interest.

## Supplementary data for

### 1 Introduction

This document describes handling of mass spectrometry data from large experiments using the `MSnbase` package and more specifically its *on-disk* backend. For demonstration purposes, the MassIVE data set MSV000080030 is used. This consists of over 1,000 mzXML files from swab-samples collected from hands and various personal objects of 80 volunteers.

### 2 Data handling and analysis with MSnbase

In this section we demonstrate data handling and access by `MSnbase` on a large experiment consisting of more than 1,000 data files.

To reproduce the analysis described in this document, download the *MSV000080030* folder from ftp://massive.ucsd.edu/MSV000080030/ and place it into the same folder as this document.

Below we load the required libraries and define the files to be analyzed.

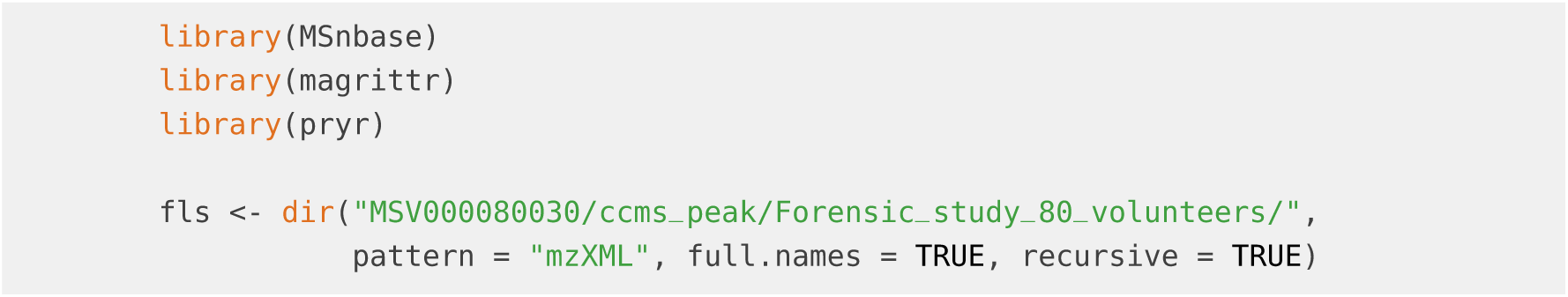

The data set consists of 1182 mzXML files. We next load the data using the two different `MSnbase` backends `“inMemory”` and `“onDisk”`. For the in-memory backend, due to the larger memory requirements, we import the data only from a subset of the files.

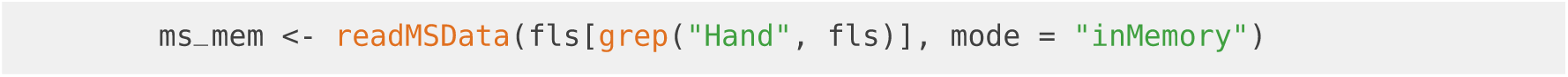

Next we load data from all mzXML files as an on-disk `MSnExp` object.

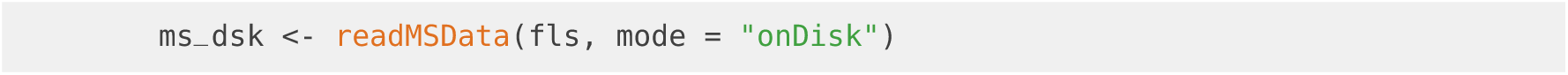

Below we count the number of spectra per MS level of the whole experiment.

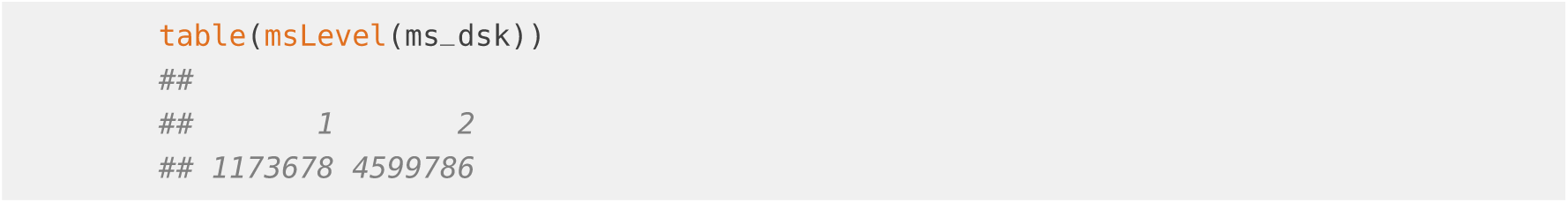

Note that the in-memory `MSnExp` object contains only MS2 spectra (in total 2140520) from a subset of data files. However, the data import was much slower (over ∼ 12 hours for the in-memory backend while creating the on-disk object from the full data data set took ∼ 3 hours).

Next we subset the on-disk object to contain the same set of spectra as the in-memory `MsnExp` and compare their memory footprint.

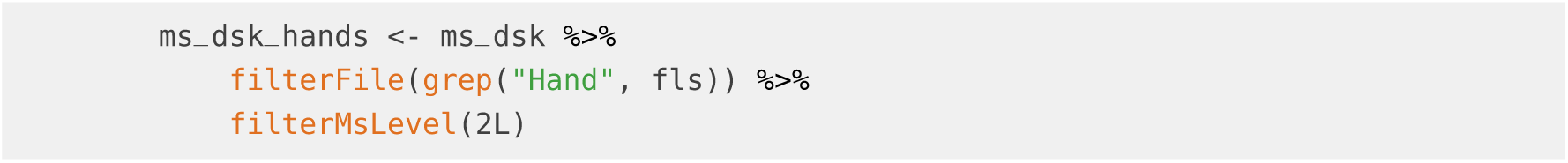

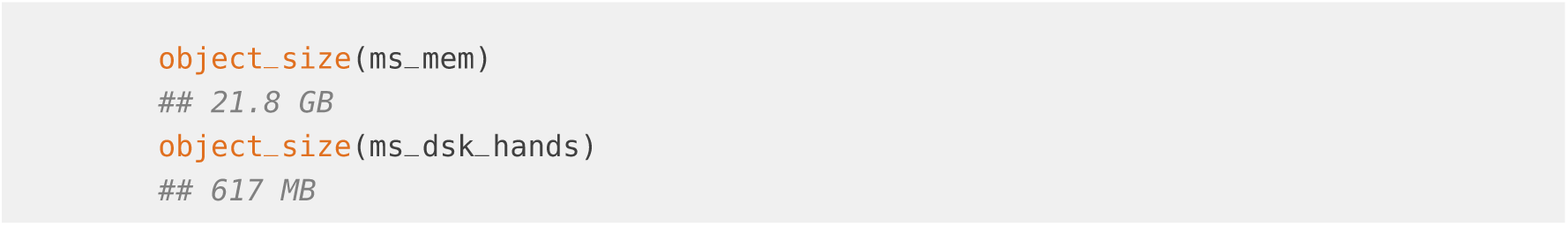

Since the on-disk object stores only spectra metadata in memory it occupies also much less system memory. As a comparison, the on-disk `MSnExp` for the full experiment was still much smaller than the in-memory object:

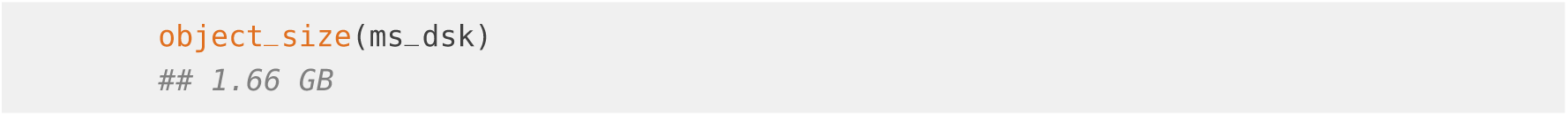

#### 2.1 Basic MS data access functionality

Before evaluating the `MSnbase` performance on the large data set we provide some general description of the `MSnbase` data classes and basic data access operations. MS data from raw data files in mzML, mzXML, mzData or netCDF format is represented by the `MSnExp` object which organizes the spectra from the original files in an one-dimensional list. Functions like `rtime` and `msLevel` allow to extract the retention time and MS level, respectively. They return a `numeric` (or `integer`) vector with the same length as the number of spectra in the `MSnExp`. In the example below we use the `rtime` function to extract the retention times for each spectrum.

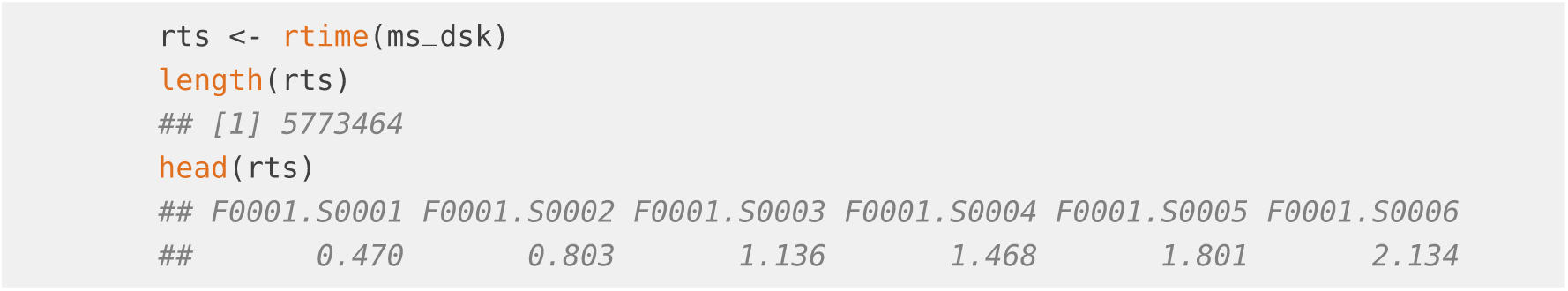

The `fromFile` function can be used to determine the source file (sample) of a specific spectrum in the `MSnExp` object. This function returns an `integer` vector, of the same length as spectra in the experiment, with the file index. The file names can be accessed with the `fileNames` method. An `MSnExp` object can be subsetted with [ and e.g. the index of the spectra that should be retained. In the code block below we subset our `ms_dsk` object to keep only spectra from the 3rd file.

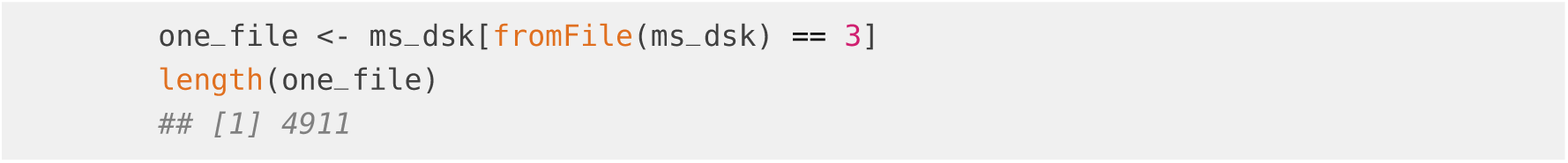

Note that there are also dedicated *filter* functions to subset an `MSnExp` object such as `filter File`, `filterMsLevel`, `filterRt`, `filterMz`, `filterPrecursorMz` or `filterIsolationWindow`. In the example below we use the `filterRt` function to further subset our data to keep only spectra acquired within a certain time range.

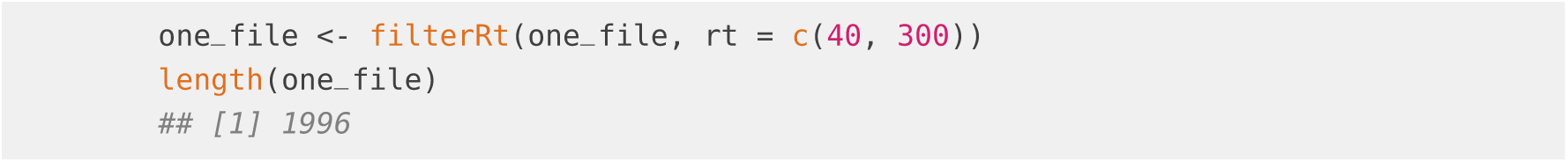

As mentioned above, the `MSnExp` object is comparable with a list of spectra. Thus, to extract a single spectrum from it we can use [[. This will return an object of type `Spectrum` which encapsules/represents all information of the measured spectrum (i.e. m/z and intensity values as well as metadata information). In the example below we extract the 15th spectrum from our data subset and access its m/z values with the `mz` function.

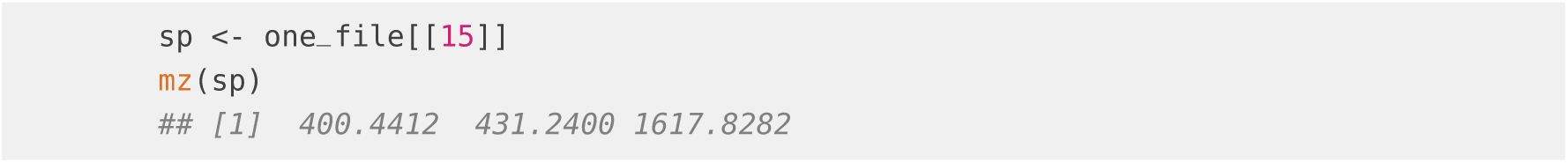

This particular spectrum has only 3 peaks.

Note that m/z or intensity values can also be directly extracted from the `MSnExp` object as shown in the example below. The result will be a `list` of `numeric` vectors, each element representing the m/z values for each spectrum in the object.

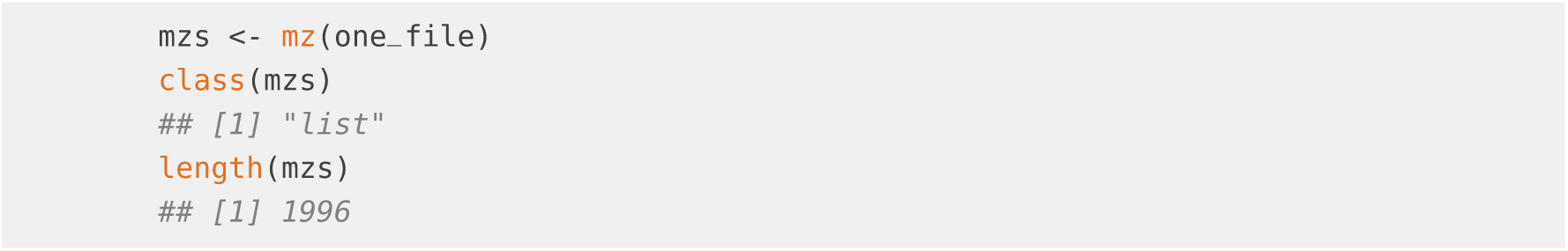

In addition, it is also possible to extract all m/z and intensity values from an `MSnExp` object as a `data.frame` as shown in the code block below, but this is not suggested, since it loads all the data into memory but all MS spectrum metadata such as MS level or precursor m/z get lost.

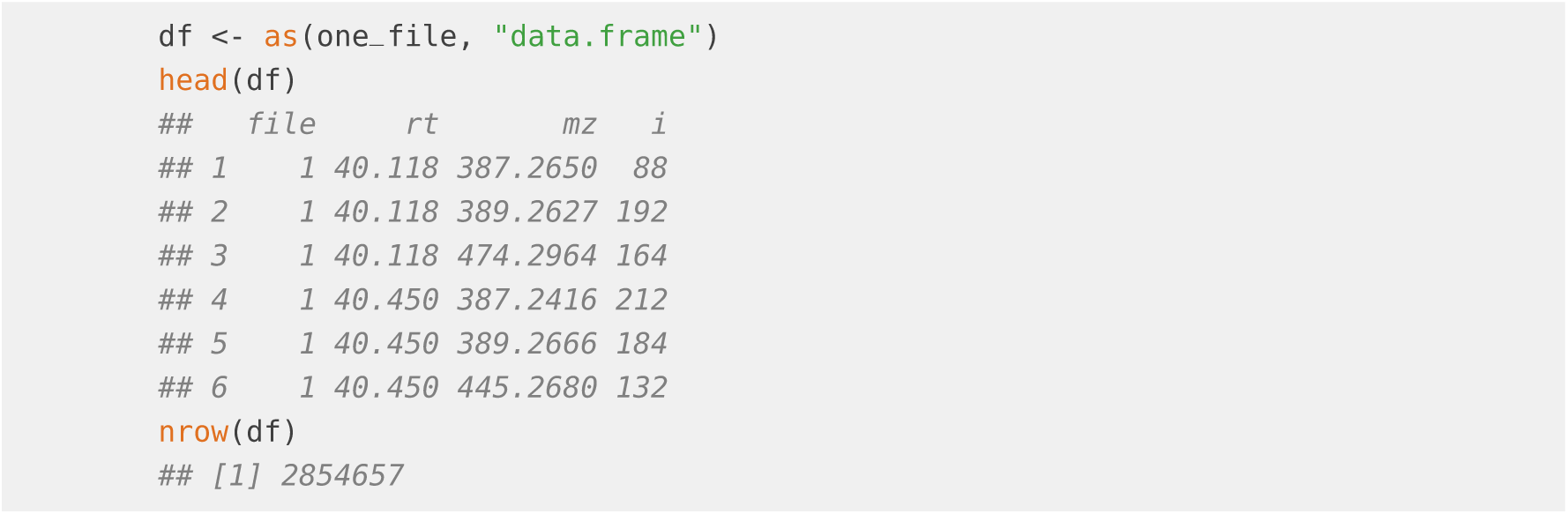

Note that for all these operations it is irrelevant whether an in-memory or on-disk backend was used. In general it is advisable to use the on-disk backend especially for experiments with more than ∼ 50 files.

#### 2.2 Performance of the on-disk backend on large scale data sets

To demonstrate `MSnbase`’s efficiency in processing large scale experiments we perform some standard subsetting, data access and manipulation operations.

We first compare the performance of the on-disk and in-memory backend on accessing m/z values with the `mz` function on a set of 100 randomly selected spectra. The performance is assessed with the `microbenchmark` function.

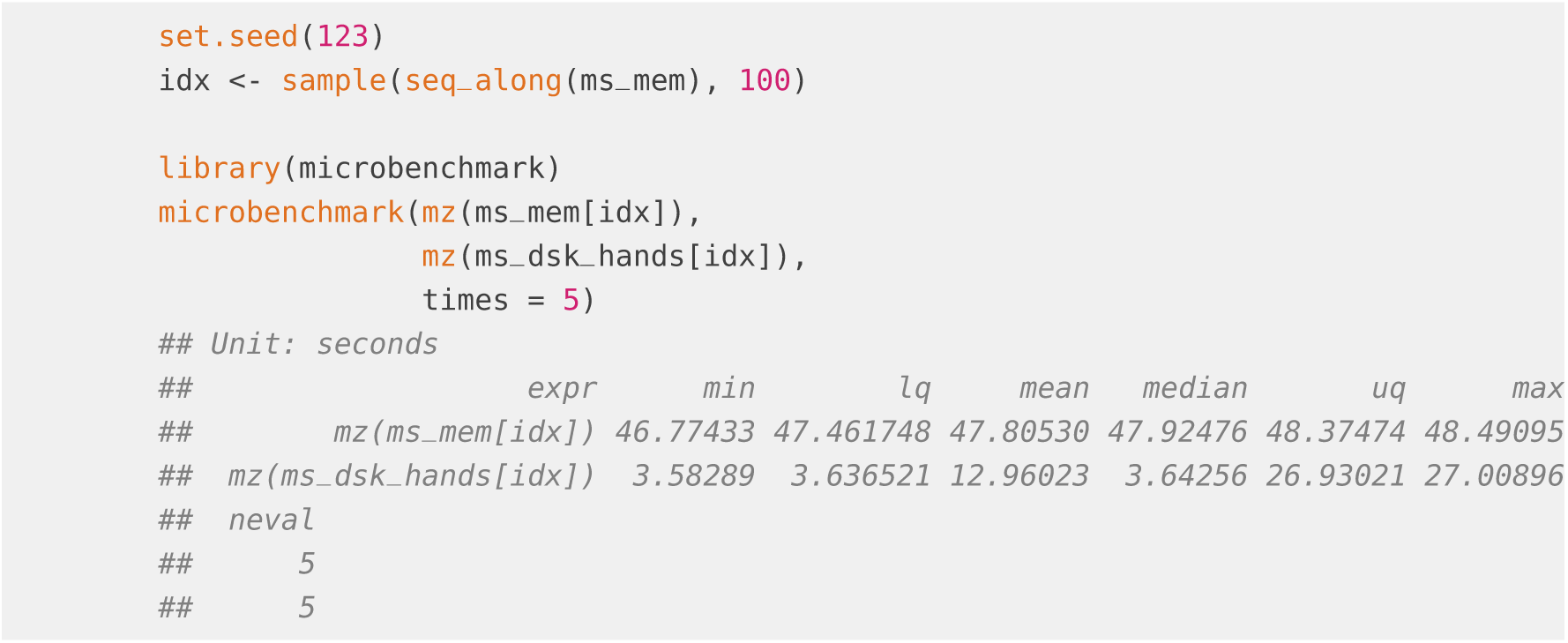

For this combined subsetting and data access operation the on-disk backend performed better than the in-memory `MSnExp`, while even requiring much less memory.

Next we extract all MS2 spectra with a retention time between 50 and 60 seconds and a precursor m/z of 108.5362 (+/- 5ppm). This subsetting operation is performed on the on-disk `MSnExp` object representing the full experiment with the 1182 data files/samples. To assess the performance of the following operations we use `system.time` calls that record elapsed time in seconds.

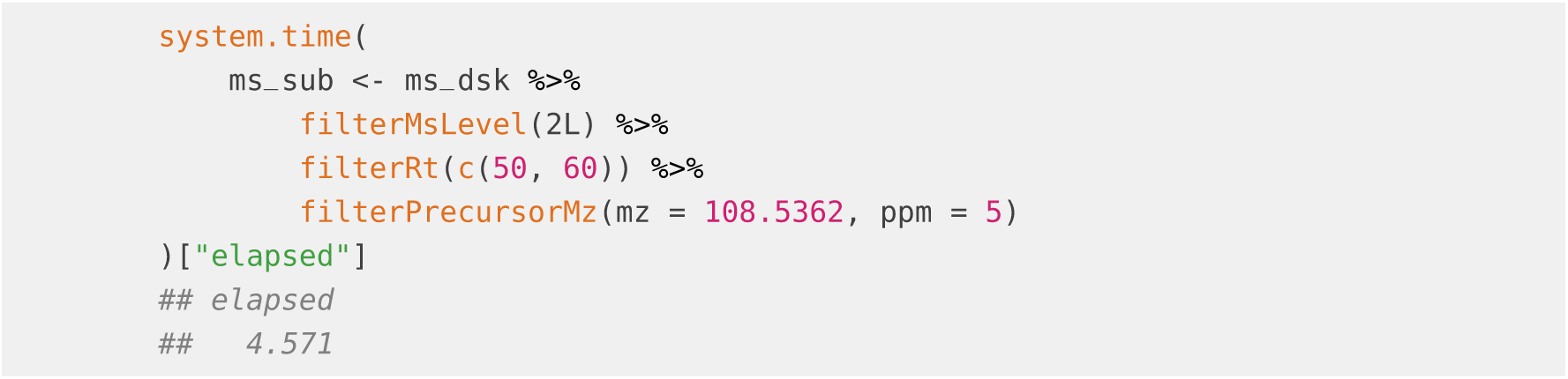

In total `length(ms_sub)` spectra were selected from in total 928 data files/samples. The plot below shows the data for the first spectrum.

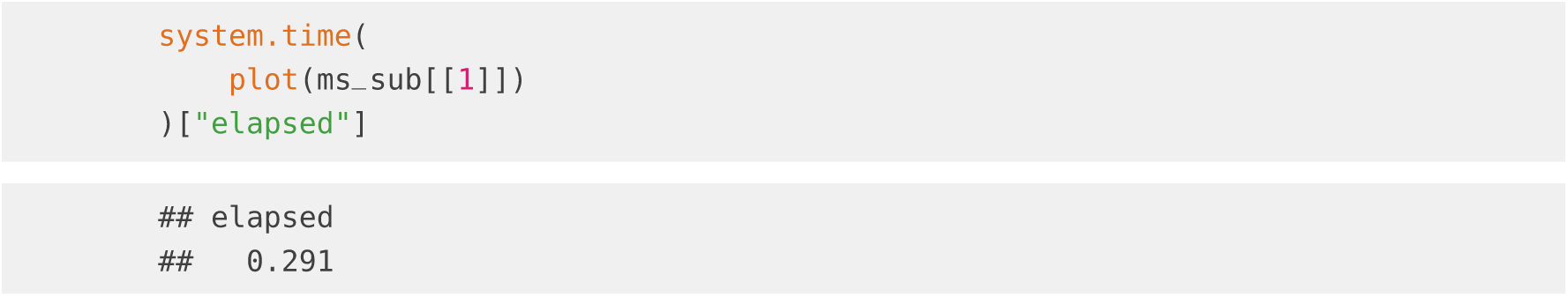

Since there seems to be quite some background noise in the MS2 spectrum we next remove peaks with an intensity below 50 by first replacing their intensities with 0 (with the `remove Peaks` call) and subsequently removing all 0-intensity peaks from each spectrum with the `clean` call. In addition we *normalize* each spectrum by dividing the maximum intensity per spectrum from the spectrum’s intensities.

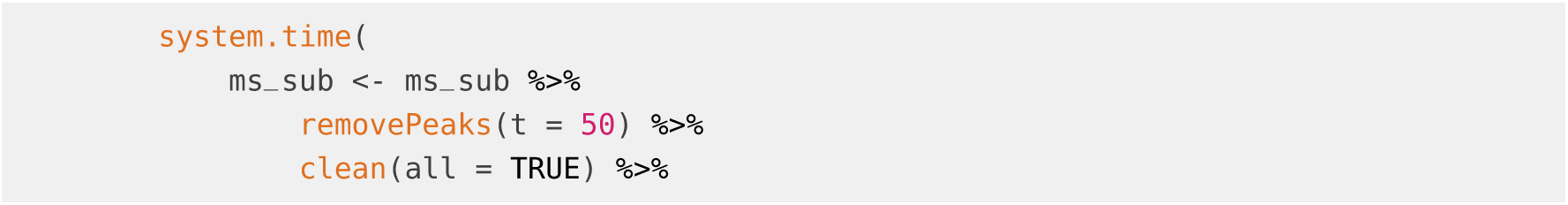

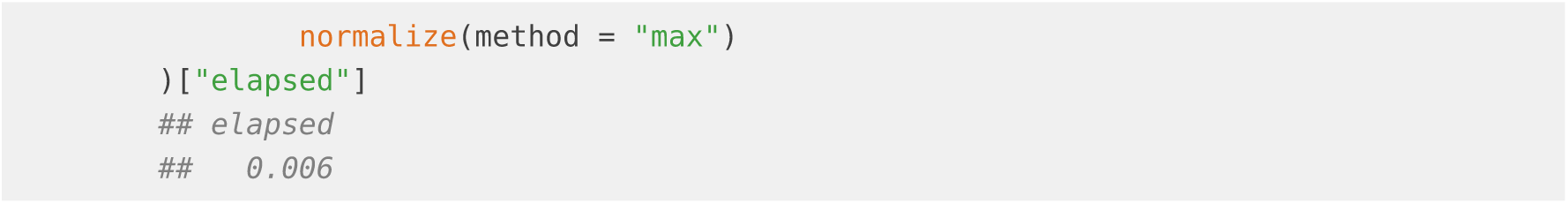

**Figure S1:**
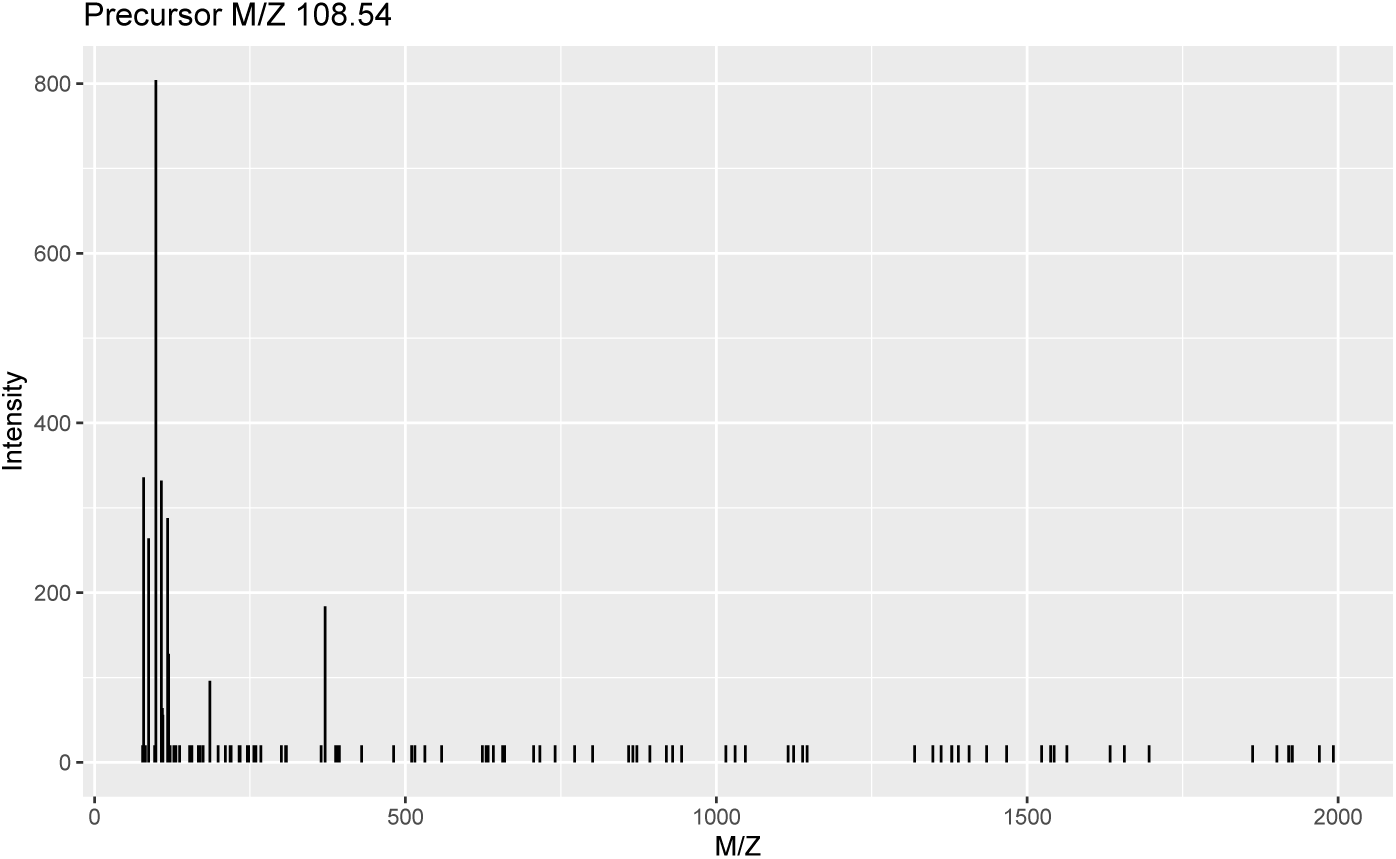
Example spectrum of the data set.

The result on the first spectrum is shown below.

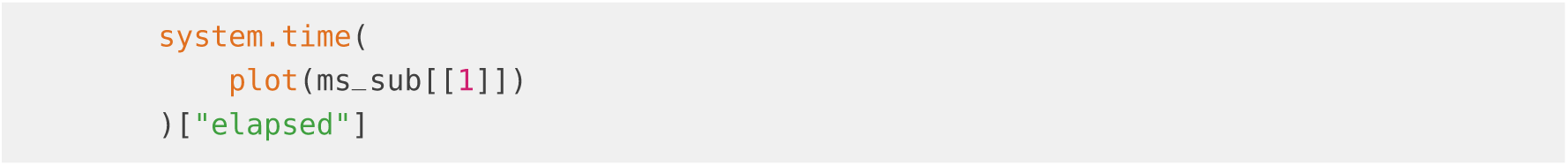

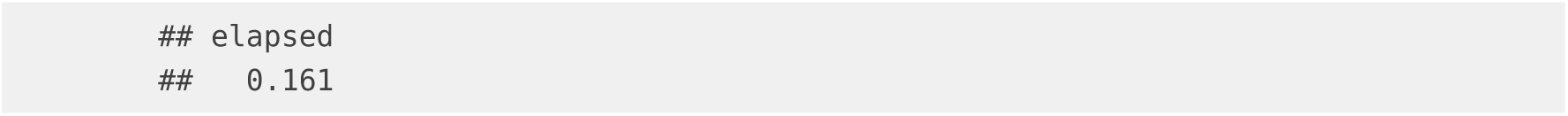

**Figure S2:**
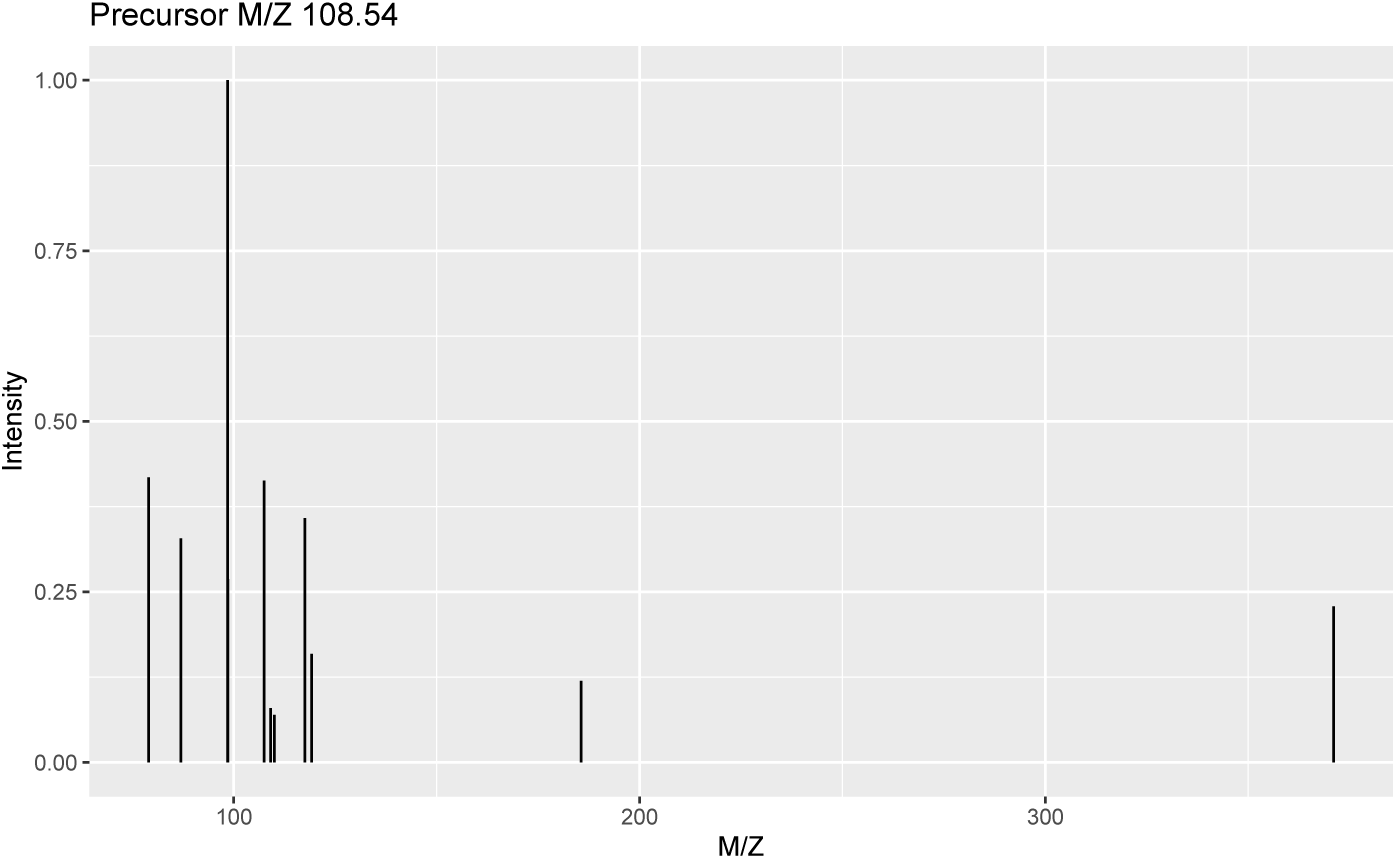
Example spectrum after cleaning.

Note that any of the data manipulations above are not directly applied to the data but *cached* in the object’s internal *lazy processing queue* (explaining the very short running time of the normalization call). The operations are only effectively applied to the data when m/z or intensity values are extracted from the object, e.g. in the `plot` call above.

For additional workflows employing `MSnbase` see also metabolomics2018^1^ that explains filtering, plotting and centroiding of profile-mode MS data with `MSnbase` and subsequent pre-processing of the (label free/untargeted) LC-MS data with the `xcms` package (that builds upon `MSnbase` for MS data representation and access).

#### 2.3 System information

The present analysis was run on a MacBook Pro 16,1 with 2.3 GHz 8-Core Intel Core i9 CPU and 64 GB 2667 MHz DDR4 memory running macOS version 10.15.5. The R version and the version of the used packages are listed below.

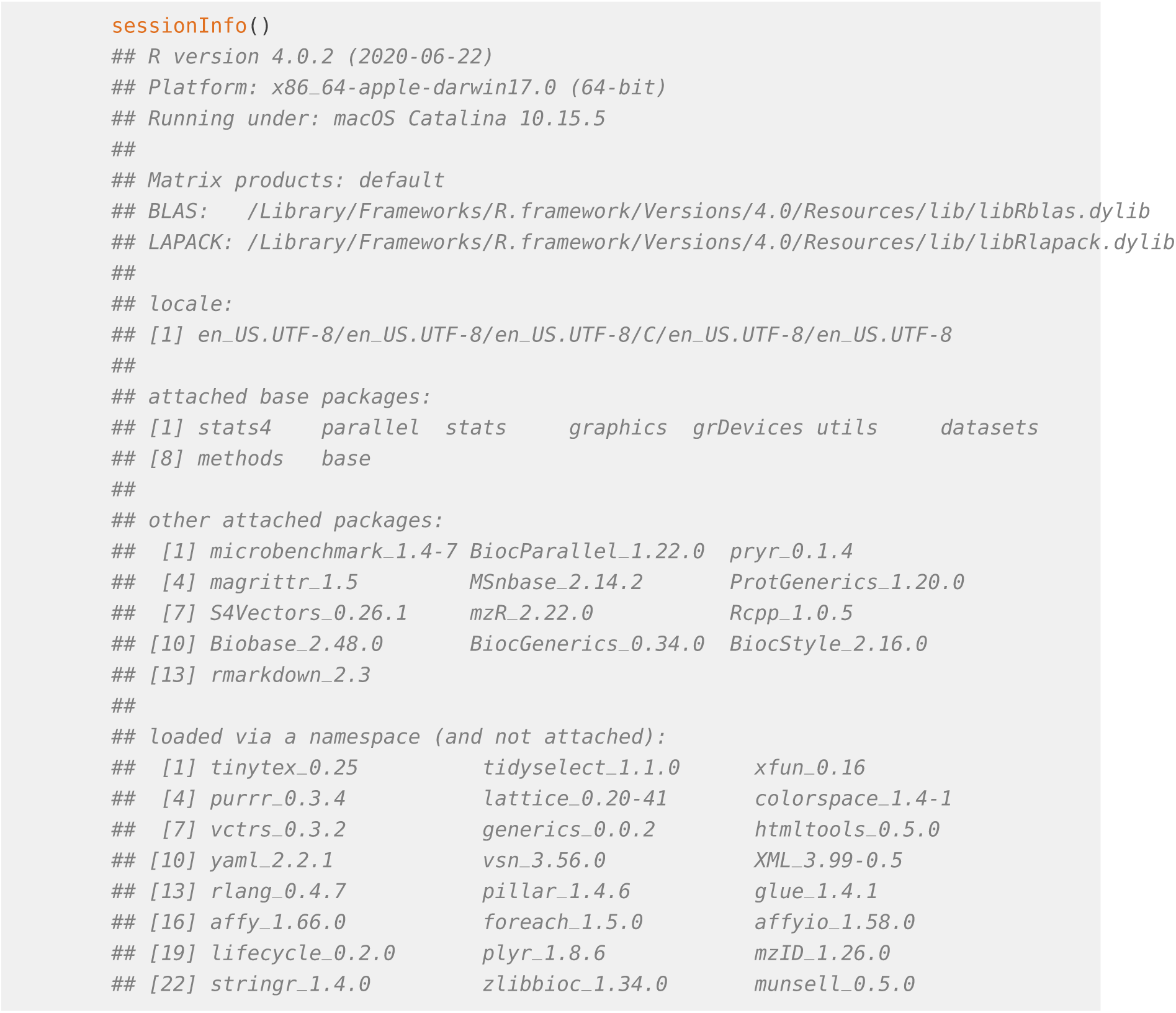

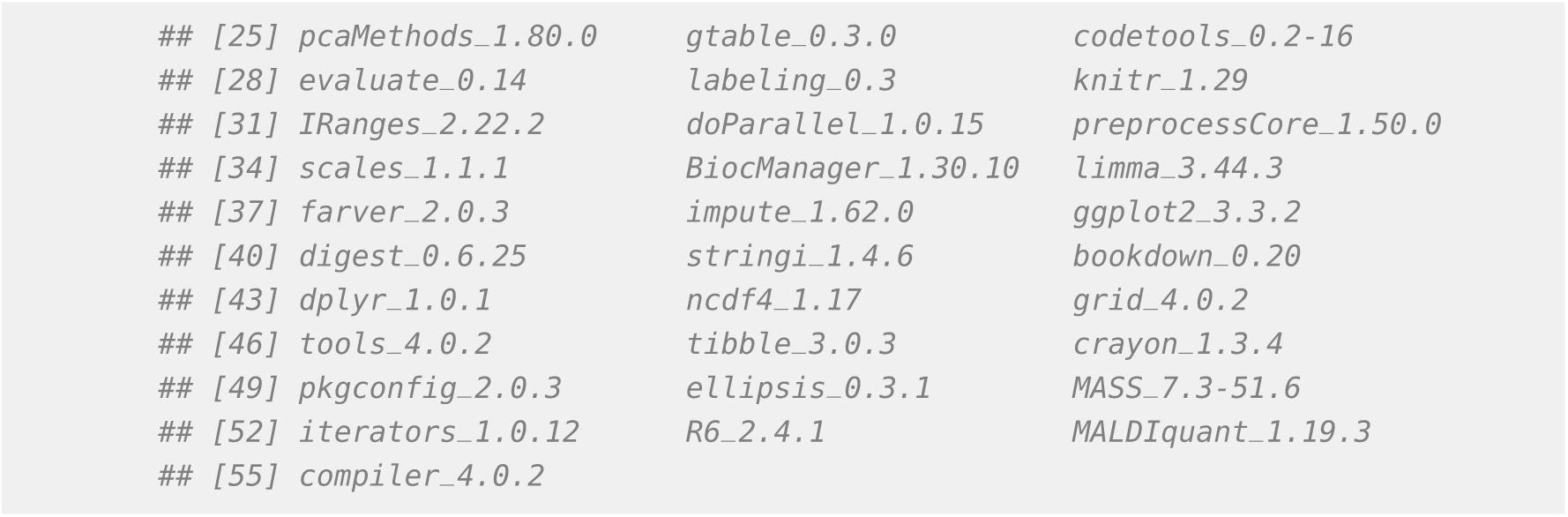

## Footnotes

1 https://github.com/jorainer/metabolomics2018

